# pyKNEEr: An image analysis workflow for open and reproducible research on femoral knee cartilage

**DOI:** 10.1101/556423

**Authors:** Serena Bonaretti, Garry E. Gold, Gary S. Beaupre

## Abstract

Transparent research in musculoskeletal imaging is fundamental to reliably investigate diseases such as knee osteoarthritis (OA), a chronic disease impairing femoral knee cartilage. To study cartilage degeneration, researchers have developed algorithms to segment femoral knee cartilage from magnetic resonance (MR) images and to measure cartilage morphology and relaxometry. The majority of these algorithms are not publicly available or require advanced programming skills to be compiled and run. However, to accelerate discoveries and findings, it is crucial to have open and reproducible workflows. We present `pyKNEEr`, a framework for open and reproducible research on femoral knee cartilage from MR images. `pyKNEEr` is written in python, uses Jupyter notebook as a user interface, and is available on GitHub with a GNU GPLv3 license. It is composed of three modules: 1) image preprocessing to standardize spatial and intensity characteristics, 2) femoral knee cartilage segmentation for intersubject, multimodal, and longitudinal acquisitions, and 3) analysis of cartilage morphology and relaxometry. Each module contains one or more Jupyter notebooks with narrative, code, visualizations, and dependencies to reproduce computational environments. `pyKNEEr` facilitates transparent image-based research of femoral knee cartilage because of its ease of installation and use, and its versatility for publication and sharing among researchers. Finally, due to its modular structure, `pyKNEEr` favors code extension and algorithm comparison. We tested our reproducible workflows with experiments that also constitute an example of transparent research with `pyKNEEr`. We provide links to executed notebooks and executable environments for immediate reproducibility of our findings.

## 1 Introduction

Open science and computational reproducibility are recent movements in the scientific community that aim to promote and encourage transparent research. They are supported by national and international funding agencies, such as the United States National Institutes of Health (NIH) [9] and the European Commission [10]. Open science refers to the free availability of data, software, and methods developed by researchers with the aim to share knowledge and tools [75]. Computational reproducibility is the ability of researchers to duplicate the results of a previous study, using the same data, software, and methods used by the original authors [5]. Openness and reproducibility are essential to researchers to assess the accuracy of scientific claims [55], build on the work of other scientists with confidence and efficiency (i.e. without “reinventing the wheel”) [54], and collaborate to improve and expand robust scientific workflows to accelerate scientific discoveries [53, 14, 42]. Historically, research data, tools, and processes were rarely openly available because of limited storage and computational power [42]. Nowadays, there are several opportunities to conduct transparent research: data repositories (e.g. Zenodo and FigShare), code repositories (e.g. GitHub, GitLab, and Bitbucket), and platforms for open science (e.g. The European Open Science Cloud and Open Science Framework). In addition, there exist computational notebooks that combine narrative text, code, and visualization of results (e.g. Jupyter notebook and R markdown), allowing researchers to create workflows that are computationally transparent and well documented [54]. Finally, it is possible to recreate executable environments from repositories to run notebooks directly in a browser and thus make code immediately reproducible (e.g. Binder).

In the evolution of research practice, the structure of scientific papers, intended as vehicles to communicate methods and results to peers, is changing. In 1992, Claerbout was among the first to envision interactive publications: “[…] an author attaches to every figure caption a pushbutton or a name tag usable to recalculate the figure from all its data, parameters, and programs. This provides a concrete definition of reproducibility in computationally oriented research” [8]. Following this vision, papers are transforming from static to interactive. They will progressively integrate data and code repositories, metadata files describing data characteristics (e.g. origin, selection criteria, etc.), and computational notebooks used to compute results and create graphs and tables [19, 21] for more transparent research.

Transparency in image-based research is crucial to provide meaningful and reliable answers to medical and biological questions [50]. In the musculoskeletal field, quantitative analysis from magnetic resonance (MR) imaging has assumed an increasingly important role in investigating osteoarthritis (OA) [22]. OA is the most common joint disease worldwide, affecting about 2 in 10 women and 1 in 10 men over 60 years of age [76]. It causes structural changes and loss of articular cartilage, with consequent pain, stiffness, and limitation of daily activities [45]. OA of the knee is one of the main forms of OA, affecting 1/3 of the adults with OA [39] and accounting for 83% of the total OA economic burden [25]. To investigate knee OA, scientists have developed algorithms to preprocess MR images, segment femoral knee cartilage, and extract quantitative measurements of morphology, such as thickness [15] and volume [56], and relaxation times, such as *T*_1*ρ*_ and *T*_2_ [32].

In the image analysis pipeline, segmentation constitute a major challenge. Researchers still tend to segment femoral knee cartilage manually or semi-automatically, using commercial or in-house software, in a tedious and non-reproducible manner [41, 35]. However, there exist several algorithms that researchers have developed to automatically segment knee cartilage. In the literature and in published reviews [24, 48, 79], we have found 29 relevant publications that propose new algorithms to segment femoral knee cartilage. These algorithms are based on different principles, namely active contours, atlas-based, graph-based, machine and deep learning, and hybrid combinations, and were developed by various research groups worldwide (Fig. 1). Of these, only the implementations by Wang et al. [69] and by Shan et al. [59] are open-source and hosted in public repositories (see Wang’s repository and Shan’s repository). These two implementations, however, have some limitations: in the first case, documentations of code and usage are not extensive, while in the second case the code is written in C++ and requires advanced programming skills to be compiled and run. Other communities, such as neuroimaging, largely benefit from robust, open-source, and easy-to-use software to segment and analyze images (e.g. ANTs [3], FreeSurfer [17], Nipype [20]). Because of these open-access tools, researchers do not need to re-implement algorithms for basic processing and can focus on further statistical analyses [65, 29, 13]. To accelerate discoveries and findings, it is fundamentally important to have not only open-source tools, but also workflows that are computationally reproducible, and thus enhance scientific rigor and transparency [14].

**Figure 1:**
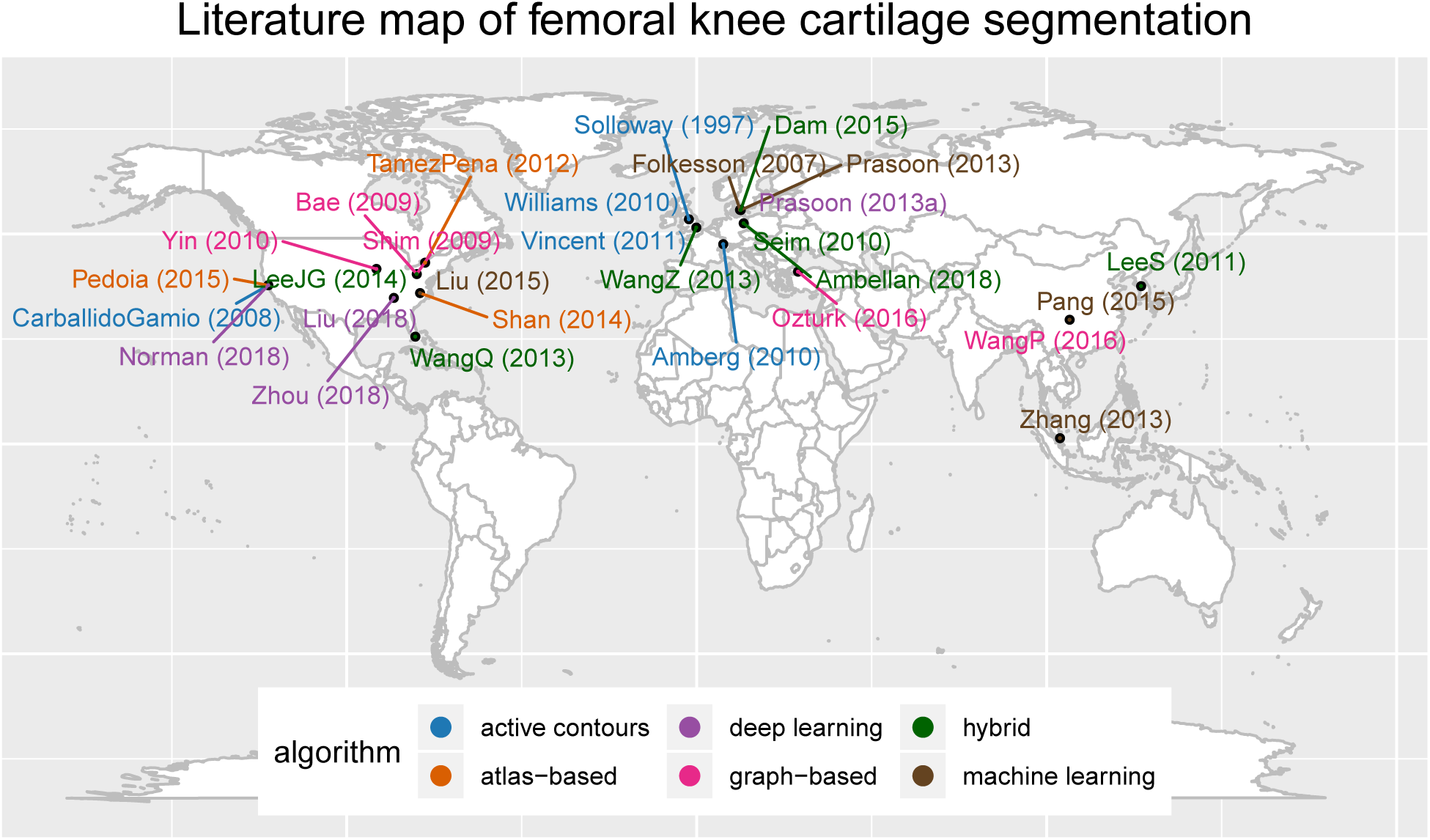
The visualization shows name of first author, year of publication, affiliation of last author, and segmentation method for 29 relevant publications on femoral knee cartilage segmentation from 1997 to 2018. Publications by segmentation method and in alphabetical order are: Active contours: Amberg(2010)[2], Carballido-Gamio(2008)[6], Solloway(1997)[62], Vincent(2011)[67], Williams(2010)[74]; Atlas-based: Pedoia(2015)[47], Shan(2014)[59], Tamez-Pena(2012)[64]; Deep-learning: Liu(2018)[33], Norman(2018)[43], Prasoon(2013a)[52], Zhou(2018)[81]; Graph-based: Bae(2009)[4], Ozturk(2016)[44], Shim(2009)[60], WangP(2016)[68], Yin(2010)[78]; Hybrid: Ambellan(2018)[1], Dam(2015)[11], LeeJG(2014)[30], LeeS(2011)[31], Seim(2010)[57], WangQ(2013)[69], WangZ(2013)[70]; Machine learning: Folkesson(2007)[18], Liu(2015)[34], Pang(2015)[46], Prasoon(2013)[51], Zhang(2013)[80]. This graph and graphs in Fig. 4 and Fig. 5 were made in Jupyter notebook using ggplot2 [71], an R package based on the grammar of graphics [72]. 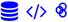

In this paper, we present `pyKNEEr`, an automatic workflow to preprocess, segment, and analyze femoral knee cartilage from MR images specifically designed for open and reproducible research. `pyKNEEr` is written in python with Jupyter notebooks as graphical user-interface, is shared on GitHub, and has a documentation website. In addition, we provide an example of transparent research with `pyKNEEr` through our validation study, implemented using images from the Osteoarthitis Initiative (OAI) [49] as well as in-house images. Finally, to be compliant with recommendations for interactive publications, throughout the paper we provide links to data files and repositories 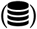, software repositories 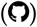, specific code and Jupyter notebooks (</>), executable environments 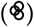, metafiles and web documentation 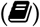, and websites [37].

## 2 Characteristics and structure of pyKNEEr

The main characteristics of `pyKNEEr` are embedded in its name: *py* is for *python*, to indicate openness, *KNEE* is for femoral knee cartilage, and *r* is for *reproducibility*.

### 2.1 Openness: python, file formats, code reuse, and GitHub

Characteristics and structure of `pyKNEEr` are based on recommendations for open scientific software in the literature, such as usage of open language and file formats, code reuse, and licensed distribution [53, 26, 54]. We wrote `pyKNEEr` in the open language python, using open-access libraries to perform computations, such as NumPy for linear algebra, pandas for data analysis, matplotlib for visualizations, SimpleITK for medical image processing and analysis [36], and itkwidgets for 3D rendering. We used widespread open-source formats for input and output data, such as text files (`.txt`) for input image lists, metafile (`.mha`) for images, and tabular files (`.csv`) for tables. To favor our code reuse, we organized `pyKNEEr` in three modules: 1) image preprocessing; 2) femoral knee cartilage segmentation; and 3) morphology and relaxometry analysis (Fig. 3). Modularity will allow us and other researchers to test, enhance, and expand the code by simply modifying, substituting, or adding Jupyter notebooks. At the same time, we reused open-source code developed by other scientists, such as preprocessing algorithms developped by Shan et al. [59] and elastix for atlas-based segmentation [28]. Finally, we released `pyKNEEr` on GitHub with a GNU GPLv3 license, which requires openness of derivative work. For citation, we assigned `pyKNEEr` a digital object identifier (DOI), obtained through the merge of the GitHub release to Zenodo (Table 1).

**Figure 2:**
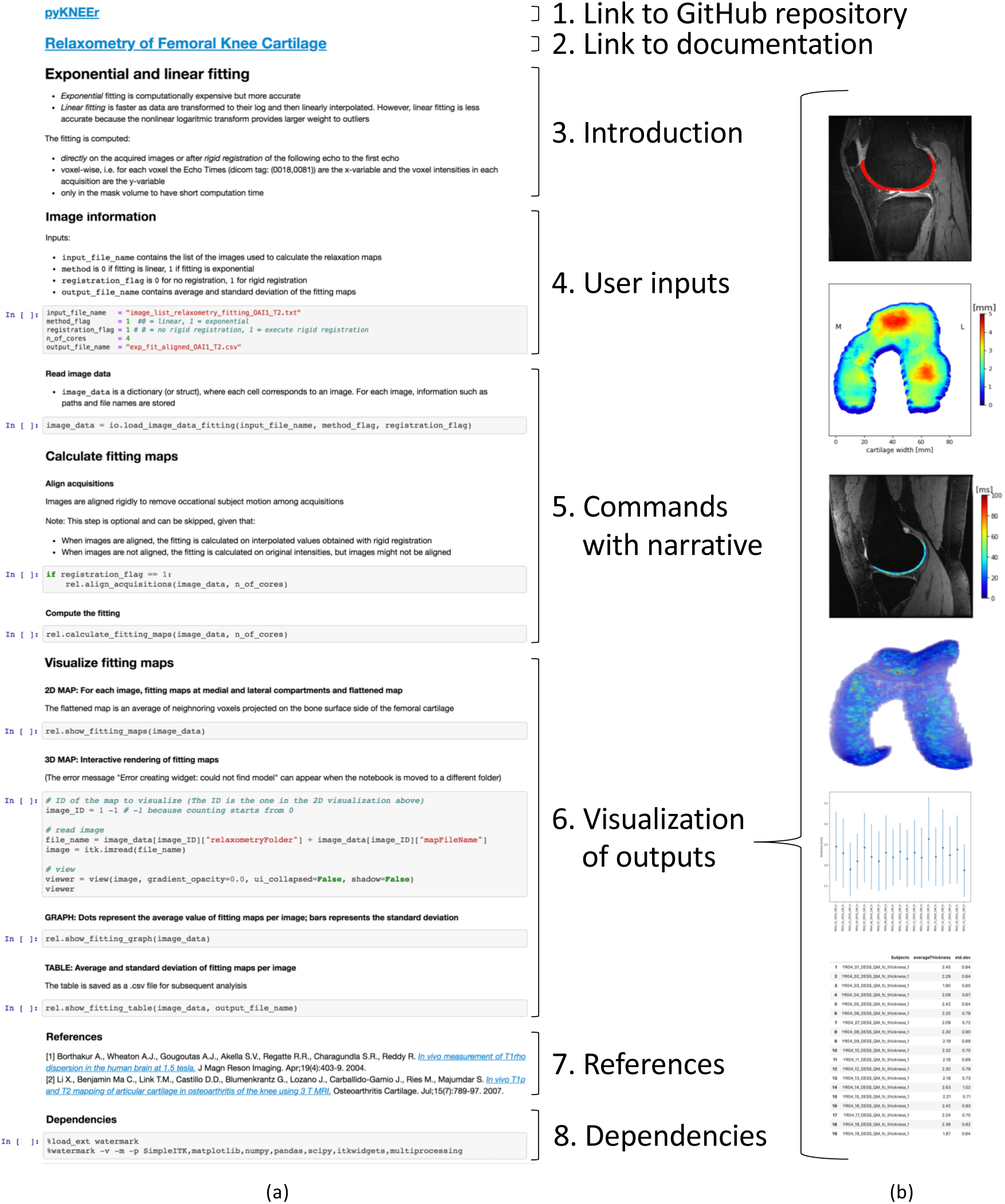
User-interface of modules in `pyKNEEr`: (a) Structure of Jupyter notebooks and (b) qualitative and quantitative visualization of outputs (from top: cartilage segmentation on image slice, flattened map of cartilage thickness, relaxation map on image slice, 3D relaxation map, and plot and table with average and standard deviation of thickness values).

**Figure 3:**
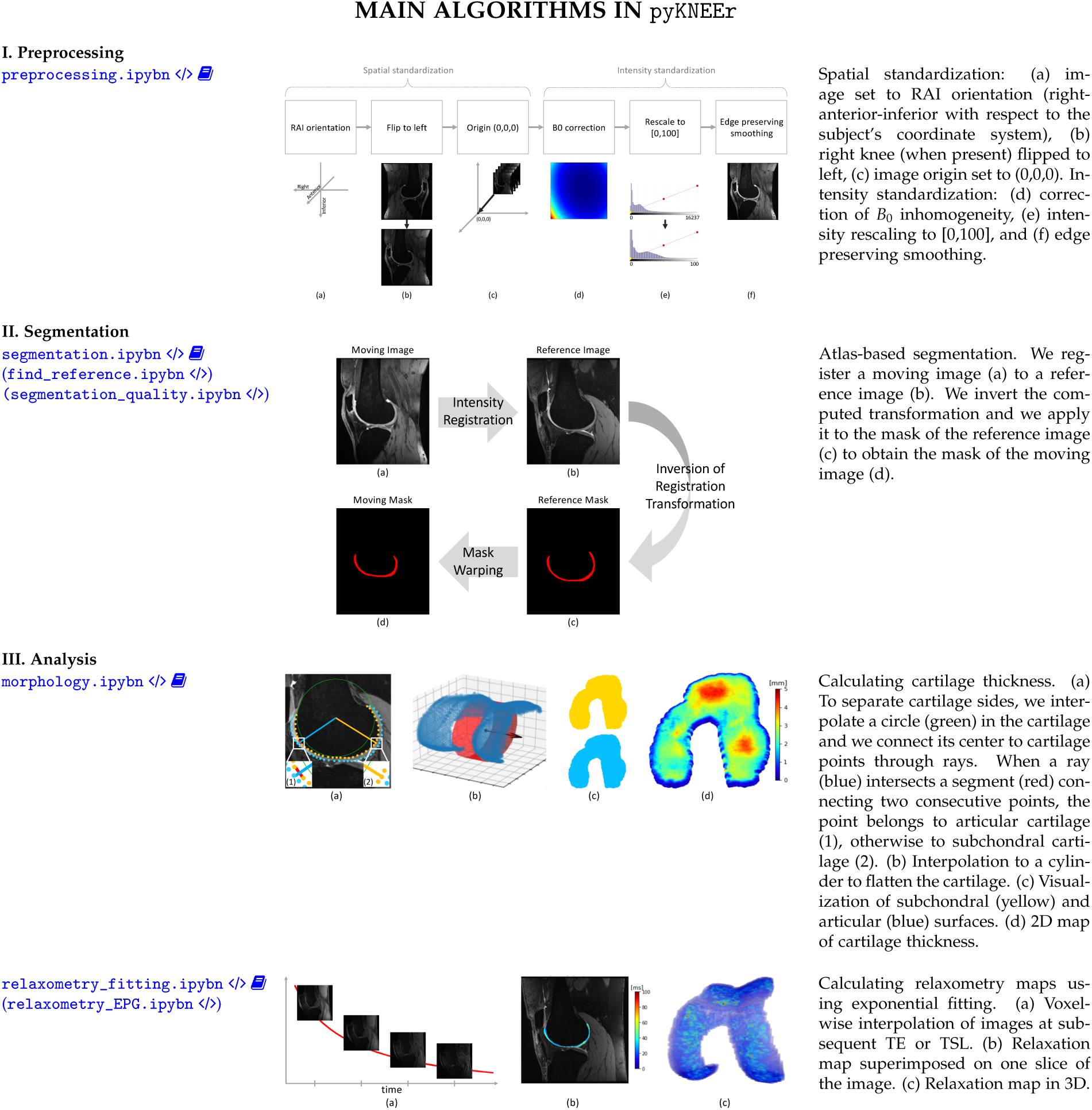
Main algorithms in `pyKNEEr` modules: I. Image preprocessing; II. Femoral cartilage segmentation; and III. Analysis of morphology and relaxometry. Left: Links to Jupyter notebooks (</>) and documentation 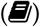. In parenthesis, notebooks in the module not depicted here. Middle: Graphic summary of algorithms. Right: Algorithm descriptions.

**Table 1:** Openness and reproducibility of `pyKNEEr` code and experimental data. DOIs and citations of used software and original data refers to their corresponding publication (*).

|  | Repository | Metadata / Documentation | Language / Format | License | DOI | Citation |
| --- | --- | --- | --- | --- | --- | --- |
| Software Used |  |  |  |  |  |  |
| Preprocessing | Bitbucket | Wiki | C++, ITK | Apache | https://doi.org/10.1016/j.media.2014.05.008* | [59]* |
| `elastix` 4.8 | GitHub | Github Wiki | C++, ITK | Apache | https://doi.org/10.1109/TMI.2009.2035616* | [28]* |
| Developed pyKNEEr | GitHub | Website | python, Jupyter notebook | GNU GPLv3 | https://doi.org/10.5281/zenodo.2574172 | Bonaretti S. et al. "pyKNEEr" (v0.0.1). Zenodo. 2019. 10.5281/zenodo.2574172 |
| Data |  |  |  |  |  |  |
| Original | OAI | Website | .dcm | Data user agreement | https://doi.org/10.1016/j.joca.2008.06.016* | [49]* |
| Derived (results) | Zenodo | Jupyter notebook | .mha, .txt, .csv | CC-BY-NC-SA | https://doi.org/10.5281/zenodo.2530608 | Bonaretti S. et al. Dataset used in (Bonaretti et al. 2019). Zenodo. 2019. 10.5281/zenodo.2530608 |

### 2.2 Reproducibility: Jupyter notebooks with computational narratives and dependencies

We designed `pyKNEEr` as a tool to perform and support computational reproducible research, using principles recommended in the literature [55, 54]. For each module of the framework, we used one or more Jupyter notebooks as a user-interface, because of their versatility in combining code, text, and visualization, and because they can be easily shared among researchers, regardless of operating systems.

Across `pyKNEEr` modules, we used the same notebook structure for consistent computational narratives (Fig. 2). Each notebook contains:

1. Link to the GitHub repository: The repository contains code and additional material, such as source files of documentation and publication;
2. Link to documentation: Each notebook is associated with a webpage containing instructions on how to create input text files, run notebooks, and evaluate outputs. Explanations include step-by-step instructions for a demo dataset, provided to the user to become familiar with the software. Single webpages are part of a documentation website, comprehensive of installation instructions and frequently asked questions. We created the website using sphinx, the python documentation generator;
3. Introduction: Brief explanation of the algorithms in the notebook;
4. User inputs: The input of each notebook is a text file (`.txt`) with folder paths and file names of images to process or analyze. Additional inputs are variables to customize the process, such as number of cores and algorithm options;
5. Commands with narrative: Titles and subtitles define narrative units of computations, and additional texts provide information about command usage and outcome interpretation. Commands in the notebook call functions in python files associated with that notebook (e.g. in the preprocessing module, the notebook `preprocessing.ipynb` calls the python file `preprocessing_for_nb.py`). In turn, associated python files call functions in core files (e.g. the python file preprocessing_for_nb.py calls `sitk_functions.py`, containing image handling functions);
6. Visualization of outputs: Qualitative visualizations include sagittal slices with superimposed cartilage mask or relaxometry map, 2D thickness maps, and 3D relaxometry maps, to allow users a preliminary evaluation of outputs. Quantitative visualizations include scatter plots and tables with numerical values and descriptive statistics (Fig. 2b), which are also saved in `.csv` files to allow researcher subsequent analysis;
7. References: List of main references used in notebook algorithms;
8. Dependencies: Code dependencies (i.e. version of python, python, python packages, and computer operating systems and hardware) to allow researchers to recreate the current computational environment and thus reproduce findings. To print dependencies, we used the python package `watermark`.

### 2.3 Algorithms in pyKNEEr

`pyKNEEr` contains specific algorithms to preprocess, segment, and analyze femoral knee cartilage from MR images. Wherever possible, we implemented multiprocessing using multiple computer cores to optimize computation effort.

#### Image preprocessing

Spatial and intensity preprocessing provide standardized high quality images to the segmentation algorithm [16]. In spatial preprocessing, we transform images to right-anterior-inferior (RAI) orientation, we flip right knees (when present) to the left laterality, and we set image origin to the origin of the cartesian system (0,0,0). In intensity preprocessing, we correct image intensities for the inhomogeneities of the static magnetic field (*B*_0_) [61], we rescale intensities to a common range [0 - 100], and we enhance cartilage edges with edge-preserving smoothing using curvature flow [58] (Fig. 3-I). Implementation of intensity preprocessing is a translation of the open access code by Shan et al. [59] 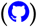 from C++ to python.

#### Femoral knee cartilage segmentation

Three steps comprise femoral cartilage segmentation: 1) finding a reference image; 2) segmenting femoral cartilage; and 3) evaluating segmentation quality (Fig. 3-II). Finding reference image and evaluating segmentation quality are possible only when ground truth segmentations are available.

##### Finding a reference image

We propose a convergence study to find the reference image, i.e. a segmented image used as a template, or atlas, for the segmentation. First, we randomly select an image as a reference image. Then, we register all images of the dataset to the reference using rigid, similarity, and spline transformations, as explained in the following paragraph. Next, we average the vector fields that result from the registrations. Finally, we choose the image whose vector field is the closest to the average vector field as the new reference for the following iteration. We repeat until two consecutive iterations converge to the same reference image or after a fixed number of iterations. It is possible to execute the search for the reference several times using different images as the initial reference to confirm the selection of the reference image. This algorithm requires femur masks because the comparison among vector fields and their average is calculated in the femur volume, as cartilage volume is limited.

##### Atlas-based segmentation

Initially, we register a moving image (i.e. any image of the dataset) to a reference image, by transforming first the femur and then the cartilage. Then, we invert the transformation describing the registration. Finally, we apply the inverted transformation to the cartilage mask of the reference image to obtain the cartilage mask of the moving image. The images to segment can be new subjects (intersubject), images of the same subject acquired at different time points (longitudinal), or images of the same subject acquired with different protocols (multimodal). To segment intersubject images, we use rigid, similarity, and spline registration, to segment longitudinal images only rigid and spline registration, and to segment multimodal images only rigid registration. We perform image registration and mask warping with elastix and transformix, respectively [28, 47], using a multiresolution approach with smoothing image pyramid, random coordinate sampler, adapitve stochastic gradient descent optimizer, and B-spline interpolators [47]. Detailed parameters are in the code repository 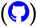.

##### Evaluating segmentation quality

We quantitatively evaluate quality of segmentation using the Dice Similarity Coefficient (DSC), a measure of the overlap between a newly segmented mask and the corresponding ground truth segmentation [12]. The Dice Similarity Coefficient is calculated as:

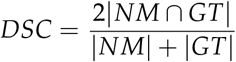

where NM is the newly segmented mask, and GT is the ground truth.

#### Morphology and relaxometry analysis

In `pyKNEEr`, cartilage analysis includes morphology and relaxometry (Fig. 3-III).

##### Cartilage morphology

Morphology quantifications are cartilage thickness and cartilage volume. To calculate cartilage thickness, first we extract contours from each slice of the cartilage mask as a point cloud. Then, we separate the subchondral side of the cartilage from the articular side, we interpolate each cartilage side to a cylinder that we unroll to flatten cartilage [41], and we calculate thickness between the two cartilage sides using a nearest neighbor algorithm in the 3D space [6, 38]. Finally, we associate thicknesses to the subchondral point cloud to visualize them as a 2D map. We compute cartilage volume as the number of voxels of the mask multiplied by the voxel volume.

##### Cartilage relaxometry

We implemented two algorithms to calculate relaxometry maps: Exponential or linear fitting and Extended Phase Graph (EPG) modeling. We use exponential or linear fitting to compute *T*_1*ρ*_ maps from *T*_1*ρ*_-weighted images and *T*_2_ maps from *T*_2_-weighted images. We calculate exponential fitting by solving a mono-exponential equation voxel-wise using a Levenberg-Marquardt fitting algorithm [32]:

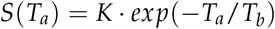

where: for *T*_1*ρ*_-weighted images, *T*_*a*_ is time of spin-lock (TSL) and *T*_*b*_ is *T*_1*ρ*_; for *T*_2_-weighted images, *T*_*a*_ is echo time (TE) and *T*_*b*_ is *T*_2_; and *K* is a constant. We compute linear fitting by transforming the images to their logarithm and then linearly interpolating voxel-by-voxel. Linear fitting is not recommended when signal-to-noise ratio is high because the logarithm transformation alters the normality of noise distribution, but it is fast and computationally inexpensive [7]. Before calculating exponential or linear fitting, the user has the option to register the images with lowest TE or TSL to the image with the highest TE or TSL to correct for image motion during acquisition [66]. We use EPG to calculate *T*_2_ maps from DESS acquisition. The implementation in `pyKNEEr` is the one proposed by Sveinsson et al. [63], which is based on a linear approximation of the relationship between the two DESS signals.

## 3 Open and reproducible research with pyKNEEr: Our validation study

We validated `pyKNEEr` with experiments that also constitute an example of open and reproducible research with `pyKNEEr`.

### 3.1 Image data

We used three datasets that we named OAI1, OAI2, and inHouse (Table 2). OAI1 contained 19 Double-Echo in Steady-State (DESS) images and *T*_2_-weighted (*T*_2_-w) spin-echo images acquired at year 4 of the OAI study. Ground truth segmentations were created using an atlas-based method (Qmetrics Technologies, Rochester, NY, USA) [64] for a previous study [23]. OAI2 consisted of 88 DESS images acquired at baseline and at 1-year followup. Ground truth segmentations were computed using an active appearance model (imorphics, Manchester, UK) [67]. Finally, inHouse contained 4 images acquired at Stanford University using DESS and CubeQuant protocols. For clarity in the following, OAI1 will be split in OAI1-DESS and OAI1-T2, OAI2 in OAI2-BL (baseline) and OAI2-FU (followup), and inHouse in inHouse-DESS and inHouse-CQ (CubeQuant). Details of the acquisition parameters are in Table 2-I.

**Table 2:**
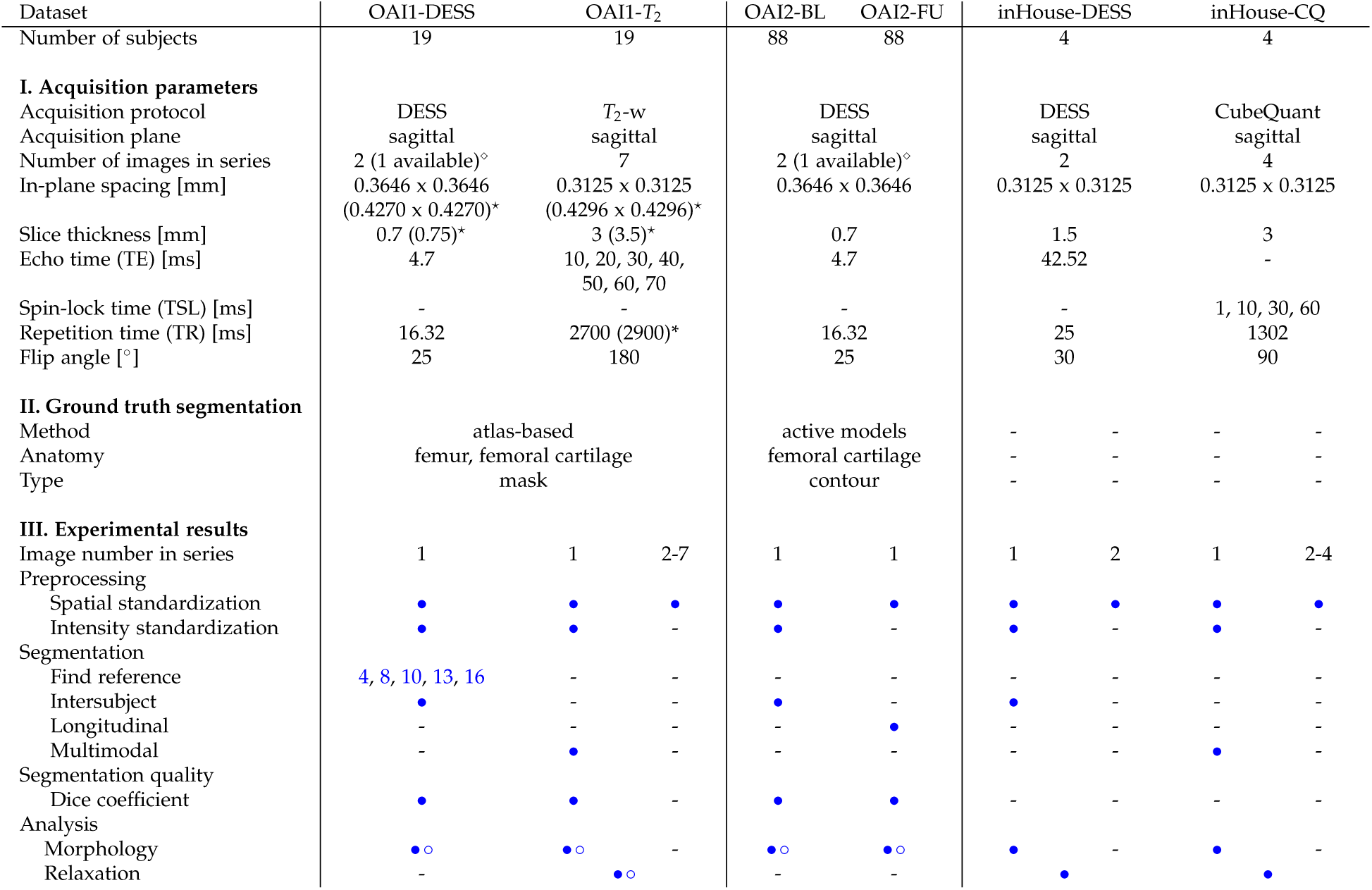
Datasets used to evaluate `pyKNEEr`. **I. Acquisition parameters**: Parameters of the protocols used to acquire the images. Images of OAI1-DESS, OAI2-BL, and OAI2-FU were acquired with the same DESS protocol, consisting of 2 echos, although only their average was available (⋄). Images of one subject of the dataset OAI1 had different slice spacing and thickness (⋆). Data queries to obtain acquisition parameters are in a Jupyter notebook 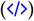. The original identification numbers (IDs) of the OAI images are in a Jupyter notebook used as a metafile 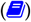. **II. Ground truth segmentation**: The datasets OAI1 and OAI2 have ground truth segmentations. They differ for computational method, segmented anatomy, and label type. **III. Experimental results**: Details of the steps in `pyKNEEr` for each dataset. Full circle (•) indicates processing of the dataset, while empty circle (∘) indicates processing of ground truth segmentations. The numbers in “Find reference” indicated the ID of the seed images used in the convergence study. Links are to the executed notebooks on GitHub.

### 3.2 Results

We preprocessed, segmented, and analyzed all the dataset using different options in `pyKNEEr`, according to dataset characteristics and availability of ground truth segmentation 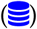 (Table 2-III).

#### Preprocessing

We executed spatial preprocessing for all images of the datasets and intensity preprocessing only for the images directly involved in segmentation.

#### Finding reference

We selected the reference mask from the dataset OAI1-DESS because of the availability of ground truth segmentations of the femur. We picked 5 images as initial reference for our parallel investigation using a python random function (random seed = 4 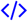). For all the studies, we found the reference as the subject whose vector field distance to the average vector field was the minimum (subject ID = 9).

#### Segmenting intersubject, longitudinal, and multimodal images

We segmented images from OAI1-DESS, OAI2-BL, and inHouse-DESS as new subjects. Segmentation failure were 1 for OAI1-DESS (ID = 6 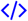), 3 for OAI2-BL (IDs = 6,24,31 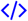), and none inHouse-DESS 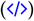. We excluded the failed registrations from the following analysis of segmentation quality, cartilage morphology, and cartilage relaxometry. We segmented the first acquisition of OAI1-T2 images 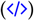 and inHouse-CQ images 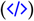 using the multimodal option in `pyKNEER`, and OAI2-FU images 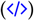 using the longitudinal option.

#### Segmentation quality

We evaluated segmentation quality for the datasets OAI1 and OAI2 because they had ground truth segmentations of femoral cartilage. The Dice similarity coefficients were 0.86 0.02 (mean standard deviation) for OAI1-DESS, 0.76 ± 0.04 for OAI1-T2, 0.73 ± 0.04 for OAI2-BL, and 0.72 ± 0.04 for OAI2-FU (Fig. 4a) 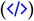.

**Figure 4:**
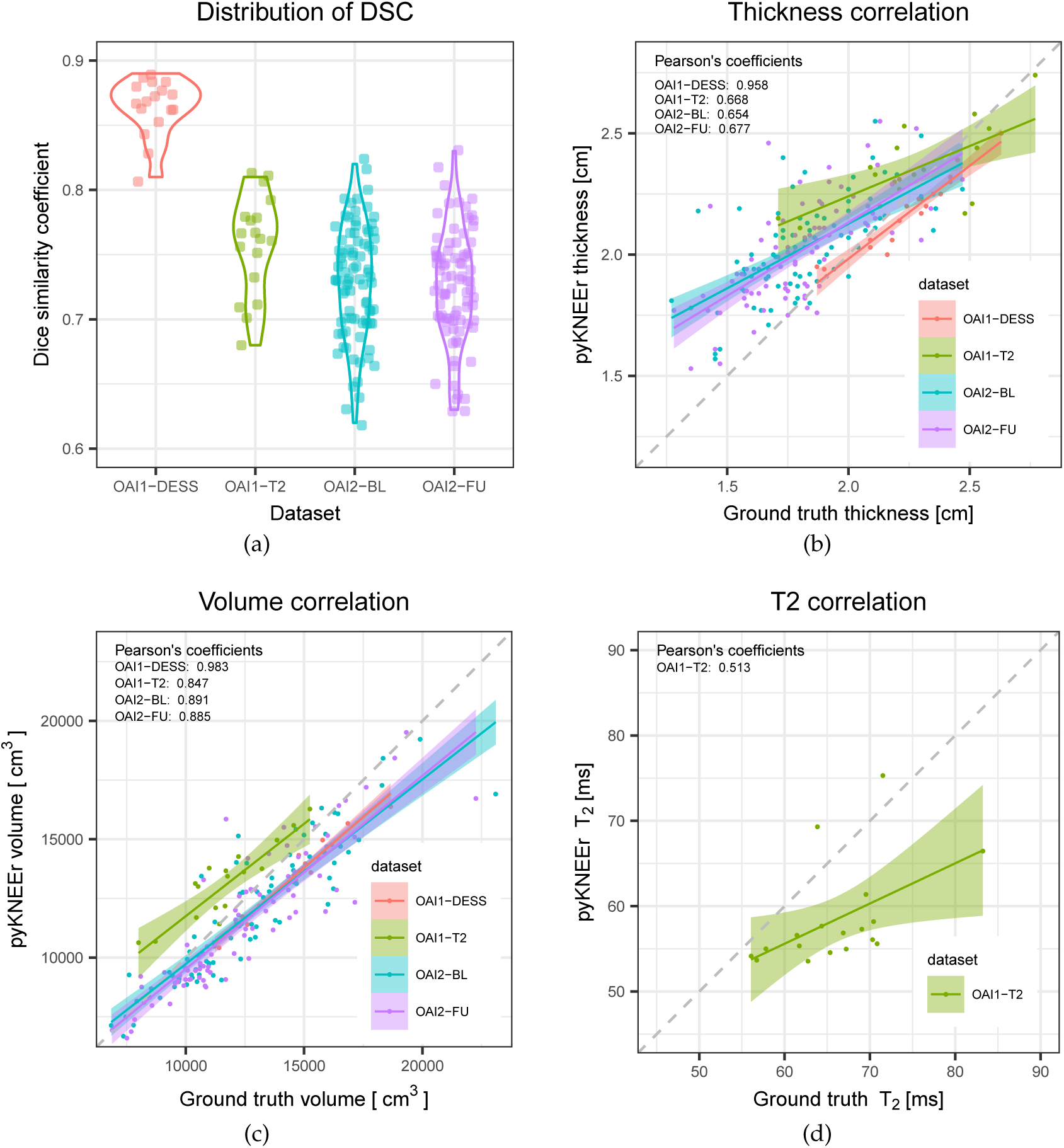
Results for the datasets OAI1-DESS (red), OAI1-T2 (green), OAI2-BL (cyan), and OAI2-FU (purple). (a) Violin plots describing the distribution of the DSC within each dataset. The dots represent DSC values spread around the y-axis to avoid visualization overlap. (b-d) Correlation between measurements derived from ground truth segmentations and `pyKNEEr`’s segmentations, i.e. cartilage thickness (b), cartilage volume (c), and *T*_2_ maps (d). 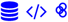

#### Morphology

We calculated cartilage thickness and volume for all datasets, including ground truth segmentations. We computed correlations of cartilage thickness calculated from `pyKNEEr`’s segmentation and ground truth segmentation, and we found that Pearson coefficients were 0.958 for OAI1-DESS, 0.668 for OAI1-T2, 0.654 for OAI2-BL, and 0.667 for OAI2-FU (Fig 4b). Similarly, we computed correlations for cartilage volume, and we found that Pearson coefficients were 0.983 for OAI1-DESS, 0.847 for OAI1-T2, 0.891 for OAI2-BL, and 0.885 for OAI2-FU (Fig 4c) 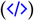.

#### Relaxometry

Before calculating relaxometry maps for OAI1-T2, we rigidly registered the images with shortest TE to the image with longest TE. Similarly, before calculating *T*_1*ρ*_ maps for inHouse-CQ, we rigidly registered the images with shortest TSL to the image with longest TSL. Then, we calculated *T*_2_ maps for OAI-T2 images extracting values in `pyKNEEr`’s masks 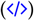 and ground truth masks 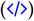, and we compared them, obtaining a Pearson’s coefficient of 0.513 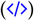. Finally, we computed relaxometry maps using exponential fitting for inHouse-CQ 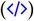 and EPG modeling for inHouse-DESS 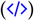 to show feasibility of the methods.

## 4 Discussion

To test possible reproducible workflows with `pyKNEEr`, we ran experiments with three different datasets. Image preprocessing was successful in all cases, while image segmentation failed in 4 cases. Average DSC were 0.81 for dataset OAI1 and 0.73 for dataset OAI2, which are in the range of published values (Fig. 5). Discrepancies of DSC between OAI1 and OAI2 can be due to the different characteristics of ground truth segmentations. OAI1 ground truth segmentations were created using an atlas-based method with *DSC* = 0.88 [64] (see “TamezPena (2012)” in Fig. 5), whereas OAI2 ground truth segmentations were created using an active appearance model with *DSC* = 0.78 [67] (see “Vincent (2011)” in Fig. 5). In addition, to calculate DSC we transformed OAI2 ground truth segmentations from contours to volumetric masks, potentially adding discretization error. Quality of segmentation had a direct impact on morphology and relaxometry analysis. Pearson’s coefficient was higher for cartilage volume than cartilage thickness, suggesting higher preservation of volume, and it was low for *T*_2_ relaxation times, suggesting higher dependency on segmentation quality for intensity-based measurements. Finally, regression lines show that measurements from `pyKNEEr` segmentation overestimated small thicknesses and underestimated large volumes and *T*_2_ values (Fig. 4). Despite its modest performance, we implemented atlas-based segmentation because it has the advantage to provide byproducts for further analysis. Image correspondences established during the registration step can be used for intersubject and longitudinal comparison of cartilage thickness and relaxation times, and voxel-based morphometry and relaxometry [47].

**Figure 5:**
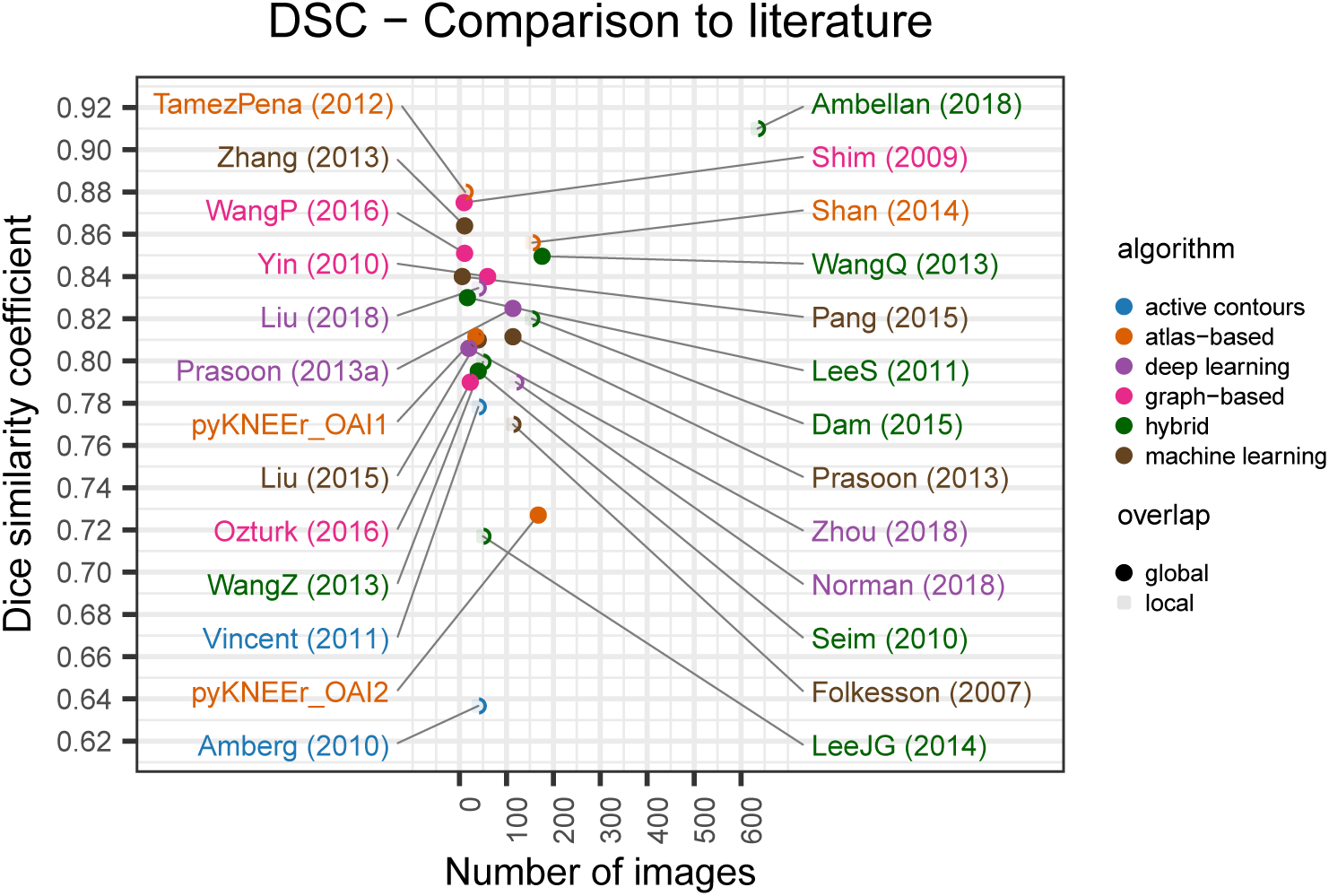
Performances of the segmentation module of `pyKNEEr`, compared with 24 studies in literature that report it. Full dots represents studies where DSCs were calculated on the whole mask, whereas empty dots represent studies where DSCs were calculated in specific parts of the cartilage, e.g. the weight-bearing area [24]. 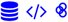

We designed `pyKNEEr` to facilitate transparent research on femoral knee analysis from MR images. Traditionally, medical image analysis workflows are in ITK, VTK, and Qt, requiring advanced computational skills in C++ to build, run, and extend code. We wrote `pyKNEEr` in python because of its ease of use, compatibility with various operating systems, and extensive computing support through packages and open code. As a consequence, `pyKNEEr` can be easily installed as a package in the python environment and does not require advanced programming skills. In addition, we used Jupyter notebooks as a user-interface because of their ease of use, literate computing approach [40], versatility for publications, and sharing among researchers. In `pyKNEEr`, Jupyter notebooks can be simply downloaded from our GitHub repository 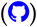 to a local folder. Researchers have to input an image list and optionally set a few variables, and after automatic execution of the notebook, they directly obtain visualizations, graphs, and tables for further analysis. In addition, researchers can link the executed notebook directly to papers (similarly to Table 2) and thus create an interactive publication with reproducible analysis. In the medical image analysis community, other examples of combined use of python and Jupyter notebooks are mainly for educational and general research purpose (e.g SimpleITK notebooks [77]), while usage of python as a programming language is rapidly gaining popularity in neuroimaging (e.g. Nipype [20]).

Several extensions of `pyKNEEr` could be imagined, due to the modularity of its structure. In the segmentation module, the current notebook implementing atlas-based segmentation (`segmentation.ipynb`) could be substituted by notebooks with hybrid machine or deep learning algorithms, which can provide higher DSC [1] (Fig.5). In the morphology module (`morphology.ipynb`), the code structure already includes a flag (`thickness_algo`) to integrate additional algorithms for cartilage thickness, such as surface normal vectors, local thickness, and potential field lines [38]. Finally, new notebooks could be added to the workflow to segment and analyze more knee tissues, such as tibial cartilage, patellar cartilage, and the menisci. Extensions will require a limited amount of effort because of the popularity and ease of python, the free availability of a large number of programming packages, and the flexibility of Jupyter notebooks [77]. In addition, standardized file format and computational environment will facilitate comparison of findings and performances of new algorithms.

In conclusion, we have presented `pyKNEEr`, an image analysis workflow for open and reproducible research on femoral knee cartilage. We validated `pyKNEEr` with three experiments, where we tested preprocessing, segmentation, and analysis. Through our validation test, we presented a possible modality of conducting open and reproducible research with `pyKNEEr`. Finally, in our paper we provide links to executed notebooks and executable environments for computational reproducibility of our results and analysis.

## Acknowledgements

This work was supported by Merit Review Award Number I01 RX001811 from the United States (U.S.) Department of Veterans Affairs Rehabilitation R%D (Rehab RD) Service and by NIHEB002524 and NI-HAR062068. The funding sources did not play a role in data analysis, manuscript preparation, or the decision to publish this work.

We would like to thank Uche Monu for her insights on cartilage segmentation, Bragi Sveinsson for sharing his code on EPG modeling, Valentina Mazzoli for her insight on relaxometry, Christof Seiler for software engineering support, Amy Silder, Julie Kolesar, and Scott Uhlrich for beta testing, Susan Holmes, Felix Ambellan, and Yolanda Gil for their feedbacks on the first preprint version of this paper, and all the researchers that make their code open-source, on whose effort we could create `pyKNEEr`.

## Conflict of interest

We do not have any conflict of interest.

## Notes

#### Summary of Updates

Updated links to GitHub for code and notebooks

https://github.com/sbonaretti/pyKNEEr

https://doi.org/10.5281/zenodo.2574172

https://doi.org/10.5281/zenodo.2530608

